# An automated microscopy workflow to study *Shigella*-neutrophil interactions and antibiotic efficacy *in vivo*

**DOI:** 10.1101/2022.09.27.509710

**Authors:** Arthur Lensen, Margarida C. Gomes, Ana Teresa López-Jiménez, Serge Mostowy

**Affiliations:** Department of Infection Biology, London School of Hygiene and Tropical Medicine, London, United Kingdom; Département de Biologie, École Normale Supérieure, PSL Université Paris, F-75005, Paris, France

**Keywords:** antibiotics, high-content microscopy, neutrophils, *Shigella*, zebrafish

## Abstract

*Shigella* are Gram-negative bacterial pathogens responsible for bacillary dysentery (also called shigellosis). The absence of a licensed vaccine and widespread emergence of antibiotic resistance has recently led the WHO to highlight *Shigella* as a priority pathogen requiring urgent attention. Several infection models have been useful to explore the *Shigella* infection process, yet we still lack information regarding events taking place *in vivo*. Here, using a *Shigella*-zebrafish infection model and high-content microscopy, we develop an automated microscopy workflow to non-invasively study fluorescently labelled bacteria and neutrophils *in vivo*. We apply our workflow to antibiotic-treated zebrafish and demonstrate that neutrophil recruitment is independent of bacterial burden. Strikingly, we discover that nalidixic acid (a bactericidal antibiotic) can restrict *Shigella* dissemination from the hindbrain ventricle. We envision that our automated microscopy workflow, applied here to study *Shigella*-neutrophil interactions and antibiotic efficacy in zebrafish, can be useful to innovate treatments for infection control in humans.

**SUMMARY STATEMENT:** We develop an automated image analysis workflow to enable fast and reliant immune cell counting in *Shigella*-infected zebrafish larvae, and reveal how antibiotics impact *Shigella*-neutrophil interactions *in vivo*.

## INTRODUCTION

*Shigella* species are the causative agent of shigellosis (or bacillary dysentery) and responsible for ∼200 000 deaths per year (Khalil et al., 2018; Kotloff et al., 2013; Kotloff et al., 2018). Among *Shigella* species, *Shigella flexneri* and *Shigella sonnei* are the most prevalent infecting humans. While *S. flexneri* mostly impacts low to middle income countries (LMICs), *S. sonnei* is more common in rich and industrialized countries (Livio et al., 2014). While no effective vaccine is currently available to prevent shigellosis (Mani et al., 2016), antibiotics are the most efficient treatment to avoid severe disease. However, the emergence of antibiotic resistance is raising significant concerns and led the World Health Organisation (WHO) to highlight *Shigella* as a priority pathogen requiring urgent attention (World Health Organization, 2017).

Considering that zebrafish larvae have no adaptive immune system, the zebrafish infection model is highly suited for studying how innate immune cells respond to *Shigella* infection (Duggan and Mostowy, 2018; Gomes and Mostowy, 2020; Mostowy et al., 2013; Torraca and Mostowy, 2018). Neutrophils are crucial to control *Shigella in vivo* (Mostowy et al., 2013; Raqib et al., 2002), but their population declines during infection due to the release of neutrophil extracellular traps (NETs) and their overall exhaustion, a condition called neutropenia. In the case of zebrafish infection with a non-lethal dose of *Shigella*, the neutrophil population can be replenished ∼48 hours post infection (hpi) by emergency granulopoiesis (Willis et al., 2018). In the case of human infection, we need to understand how antibiotics may impact *Shigella*-neutrophil interactions *in vivo* to develop better therapeutic strategies.

In this study, we exploit the zebrafish infection model and imaging power of high-content microscopy to investigate how antibiotics work in combination with neutrophils to combat *Shigella* infection *in vivo*. We first develop a workflow to automate image analysis capturing bacterial burden and neutrophil dynamics in zebrafish over time. Using our automated microscopy workflow, we test 4 different antibiotics on *Shigella*-infected zebrafish and show how they impact bacterial burden, neutrophil recruitment and bacterial dissemination from the hindbrain ventricle (HBV). These results highlight the importance of testing antibiotic efficacy *in vivo*, and illuminate a powerful approach to perform high-throughput drug screening on infected zebrafish to innovate treatments for infection control in humans.

## RESULTS

### Non-invasive characterization of bacterial burden *in vivo* over time

To quantify bacterial burden from infected zebrafish, larvae are classically dissociated in 0.1% Triton, serially diluted and spread on a Tryptic Soy Agar (TSA) plate to later count colony forming units (CFUs). However, this method is invasive and cannot be used to analyse the same larva over time. To non-invasively quantify bacterial burden from infected larvae over time, we used high-content microscopy to image up to 96 larvae per session. We designed an ImageJ macro to calculate total fluorescence in the zebrafish HBV infected with GFP expressing *S. flexneri* M90T (**Fig. 1A**), and total HBV fluorescence was compared to the number of CFUs manually quantified from the same larva. This automated microscopy workflow was performed on larvae at 2 days post fertilisation (dpf) injected with a low (10 000 CFU) or high (20 000 CFU) input of *S. flexneri* and imaged at 4 and 24 hours post-infection (hpi), to provide a broad range of infection conditions and bacterial burdens. Automated analysis of total HBV fluorescence is weakly correlated to results experimentally obtained by counting CFUs, with an R^2^ of ∼0.50 for both low or high bacterial burden (**Fig. 1B, 1C**). Results obtained in larvae having a lower bacterial burden have a smaller confidence interval (and thus a greater accuracy) than results obtained in highly infected larvae, independently of the observed timepoint. Consistent with this, the range of CFUs associated to total HBV fluorescence is also smaller, suggesting that automatic quantifications in larvae with a low bacterial burden are more precise than larvae with a high bacterial burden (**Fig. 1D)**. We therefore chose to differentiate between low and high burden when predicting *S. flexneri* burden based on total HBV fluorescence. Although automated image analysis failed to provide precise CFU quantifications for bacterial inputs tested here, it can enable qualitative description of bacterial burden (e.g. infer bacterial replication or clearance) in a non-invasive manner.

**Figure 1:**
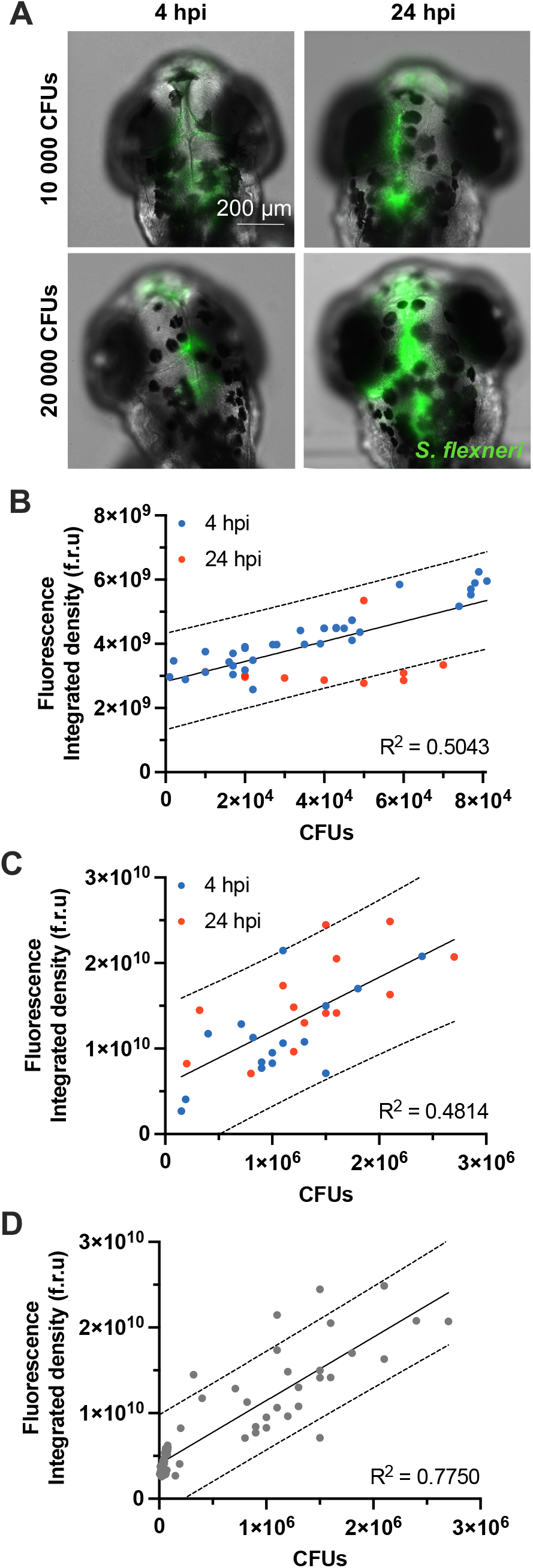
Quantification of bacterial burden in the HBV. All data presented here is collected in the HBVs of *S. flexneri* M90T-infected 2 dpf zebrafish larvae. (A) Representative images of infected larvae injected with 10 000 or 20 000 CFUs at 4 and 24 hpi. Images are maximum intensity Z-projections. Scale bar 200 μm. Green: *S. flexneri* M90T. Graphs (B-D) represent the total fluorescence in the Z-stack image of M90T-infected zebrafish larvae HBVs, at different infectious doses, measured at 4 (blue dots) or 24 hpi (red dots), and correlated to the precise number of CFUs (experimentally assessed). (B) Correlation between total fluorescence and precise number of CFUs in larvae with low bacterial burden, n=45. (C) Correlation between total fluorescence and precise number of CFUs in larvae with high bacterial burden, n=45. (D) Illustration of the challenge to accurately predict both low and high bacterial burden simultaneously, n=90. Full dark lines: linear regressions. Dashed lines: 95% confidence intervals.

Considering these limitations, we hypothesized that dark melanophores in the HBV of the developing larvae could partially block acquisition of fluorescence. To test this, we treated larvae with phenylthiourea (PTU) to inhibit melanisation (Karlsson et al., 2001), injected them with low or high input of *S. flexneri* for imaging at 4 and 24 hpi. In this case, automated analysis showed that the linear correlation between total HBV fluorescence is not improved in transparent PTU-treated larvae (**Fig. S1**).

### Use of automated microscopy to quantify zebrafish neutrophils *in vivo*

To characterize interactions between *S. flexneri* and zebrafish neutrophils, we infected larvae with neutrophils expressing the fluorescent protein DsRed (Tg(*lyz*::DsRed)^*nz50*^) and imaged them from 2 to 24 hpi using high-throughput microscopy. Although imaging of neutrophils is relatively fast, the time required to manually analyse images (∼1.5 min per image in our tested conditions) is a significant limitation. To rapidly detect neutrophils without user bias, automated image analysis workflows are required. A first method, hereafter referred to as the ‘Ellett and Lieschke’ method, uses an area calculation and averaging workflow to return a proxy of neutrophil number per image (Ellett and Lieschke, 2012). However, this method offers relatively slow analysis time (∼1.0 min per image) and imprecise neutrophil counts (ranging from 35% to 75% error rates in our tested conditions, **Fig. S2**). This imprecision is particularly apparent when analysing images of zebrafish undergoing emergency granulopoiesis, where neutrophils are densely packed together in the aorta-gonad-mesonephros (AGM, **Fig. 2A**).

**Figure 2:**
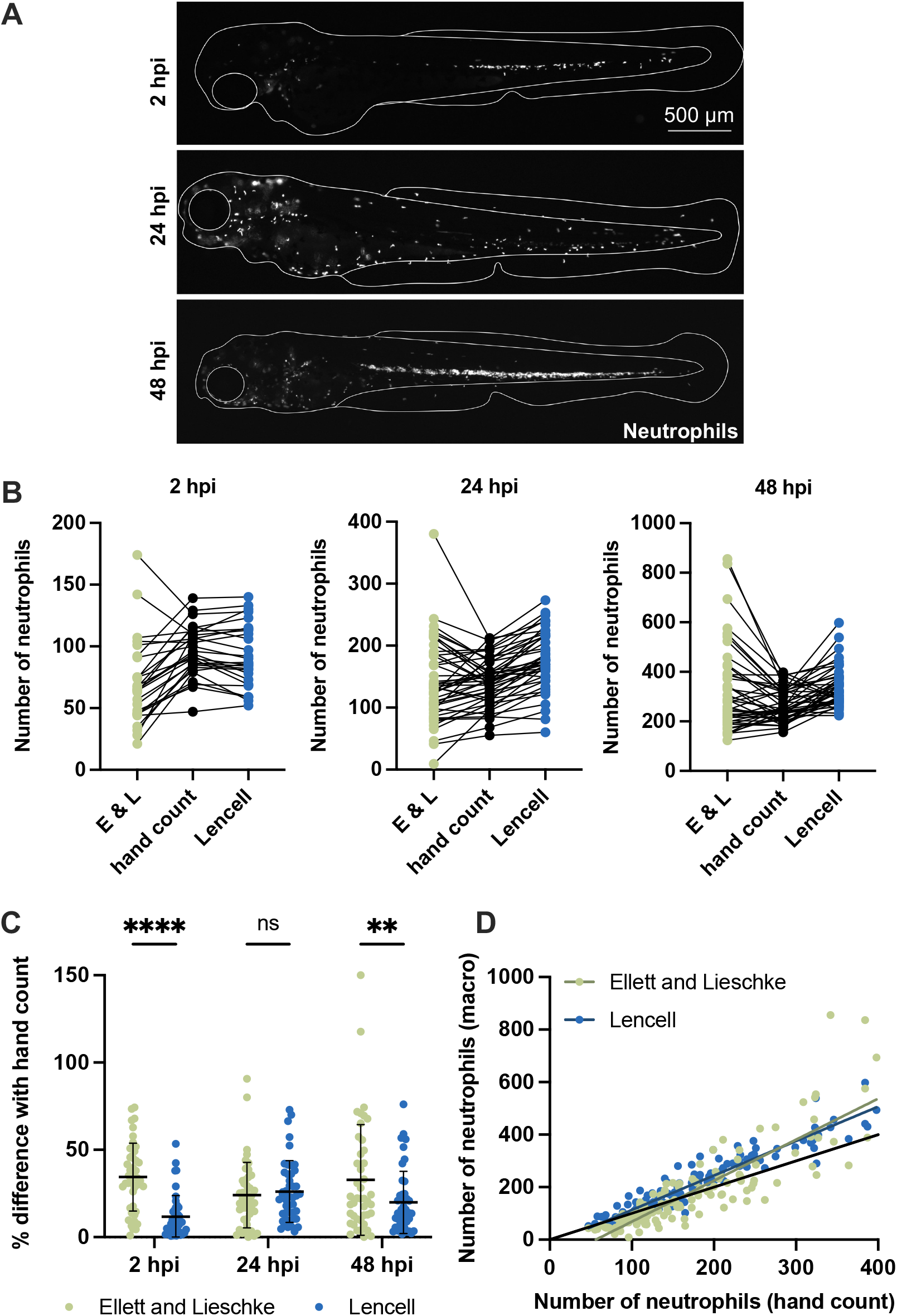
Quantification of zebrafish neutrophils. All data presented here is collected from *S. flexneri* M90T-infected 2 dpf zebrafish larvae (2 500 CFUs) at 2, 24 and 48 hpi, or PBS injected larvae. (A) Representative images of infected Tg(*lyz*::DsRed)^*nz50*^ larvae. Neutrophil counting in larvae at 48 hpi (during emergency granulopoiesis) is particularly challenging. Scale bar: 500 μm. White: neutrophils. (B) Comparison between hand counted neutrophils (black), the Ellett and Lieschke method (green) and the LenCell method (blue) in infected larvae at 2, 24 and 48 hpi, n=44 per method. (C) Comparison between hand counted neutrophils and the Ellett and Lieschke method (green) or the LenCell method (blue) in infected larvae at 2, 24 and 48 hpi. The values plotted are the differences in percentage between the macro count and the hand count, for every sample, n=44 per method (**p < 0.01, ****p < 0.0001. Two-way ANOVA with Sidak’s multiple comparison test). Black bars: mean ± SEM. (D) Correlation between hand count (x-axis) and the Ellet and Lieschke method count (green) or the LenCell method (blue) (y-axis), n = 132 per method. Black line: identity line. Green line: linear regression on the Ellett and Lieschke method results, slope = 1.561. Blue line: linear regression on the LenCell method results, slope = 1.312.

To more rapidly and precisely quantify zebrafish neutrophils from large datasets, we developed an ImageJ macro called ‘LenCell’. LenCell automatically applies a band-pass filter to remove high and low background frequencies, and to detect bright spots (in this case fluorescent neutrophils) in the cleaned images (see Methods). To characterize the efficiency of LenCell, transgenic zebrafish larvae with DsRed-neutrophils were imaged during a non-lethal *S. flexneri* infection (2500 CFU, **Fig. 2A**). We then compared quantifications performed manually, using the Ellett and Lieschke method, or using LenCell (**Fig. 2B**). By determining the mean error between the three methods from 132 images of *S. flexneri* infected larvae, we showed that LenCell is significantly more accurate than the Ellett and Lieschke method (**Fig. 2C**). Although the difference between automated quantification methods for zebrafish undergoing neutropenia (24 hpi) is negligible, LenCell is significantly more accurate when the immune system is at homeostasis (2 hpi) as well as when it is undergoing emergency granulopoiesis (48 hpi, **Fig. 2C**). As one limitation, LenCell may overestimate the number of neutrophils in larvae presenting high neutrophil numbers (> 250, **Fig. 2D**).

Overall, these results show that LenCell is significantly more accurate than the Ellett and Lieschke method at quantifying leukocytes in zebrafish larvae. LenCell is also 60x faster, as it can automatically analyse around 60 images in ∼1.0 min.

### Antibiotics reduce bacterial burden and neutrophil recruitment

To investigate the interplay between antibiotics and neutrophils during *Shigella* infection, we selected four antibiotics used clinically to treat shigellosis in humans: Nalidixic Acid (NAL), Chloramphenicol (CM), Azithromycin (AZI) and Trimethoprim (TPI). These antibiotics have different mechanisms of action in that NAL and TPI are bactericidal whereas CM and AZI are bacteriostatic (https://go.drugbank.com). TPI, CM and AZI have been shown to affect neutrophil function in humans: TPI can trigger neutropenia (De Manzini et al., 1990), whereas CM and AZI can reduce the release of neutrophil extracellular traps (NETs) (Bystrzycka et al., 2017); CM can additionally induce oxidative stress in neutrophils (Páez et al., 2008).

Considering differences in susceptibility to antibiotics between *S. flexneri* and *S. sonnei* (CDC, 2010; EUCAST, 2022), we performed zebrafish infection experiments using both bacterial species. We hypothesized that their differences in antibiotic susceptibility trigger a range of different behaviours in antibiotic-treated zebrafish, which may provide more information on the role of antibiotic-neutrophil interactions during *Shigella* infection. We first determined the minimum inhibitory concentrations (MICs) of drugs against *S. flexneri* and *S. sonnei in vitro* (**Table S1**), and observed that *S. sonnei* is less sensitive to antibiotics than *S. flexneri*. Therefore, for experiments performed *in vivo*, we used *S. sonnei* MICs because they are effective against both *Shigella* species and have no effect on larval development (data not shown).

To decipher the interplay between antibiotics, bacterial burden and neutrophil response, infections were performed in 2 dpf zebrafish larvae with DsRed labelled neutrophils. Larvae were injected in the HBV with a low input (10 000 CFU) of *S. flexneri* (**Fig. 3**) or *S. sonnei* (**Fig. S3**), and imaged every 2 h for 24 hpi using high-content microscopy. Consistent with expectations, bacterial burden is significantly reduced over time in the presence of antibiotics (**Fig. 3A**). Although the concentration of antibiotics used is 2x the MIC for *S. sonnei*, it is not enough to completely inhibit bacterial growth, indicating that MICs *in vitro* are lower than MICs *in vivo*. In agreement, we failed to capture a significant reduction of *S. sonnei* in the presence of antibiotics *in vivo* (**Fig S3A**).

**Figure 3:**
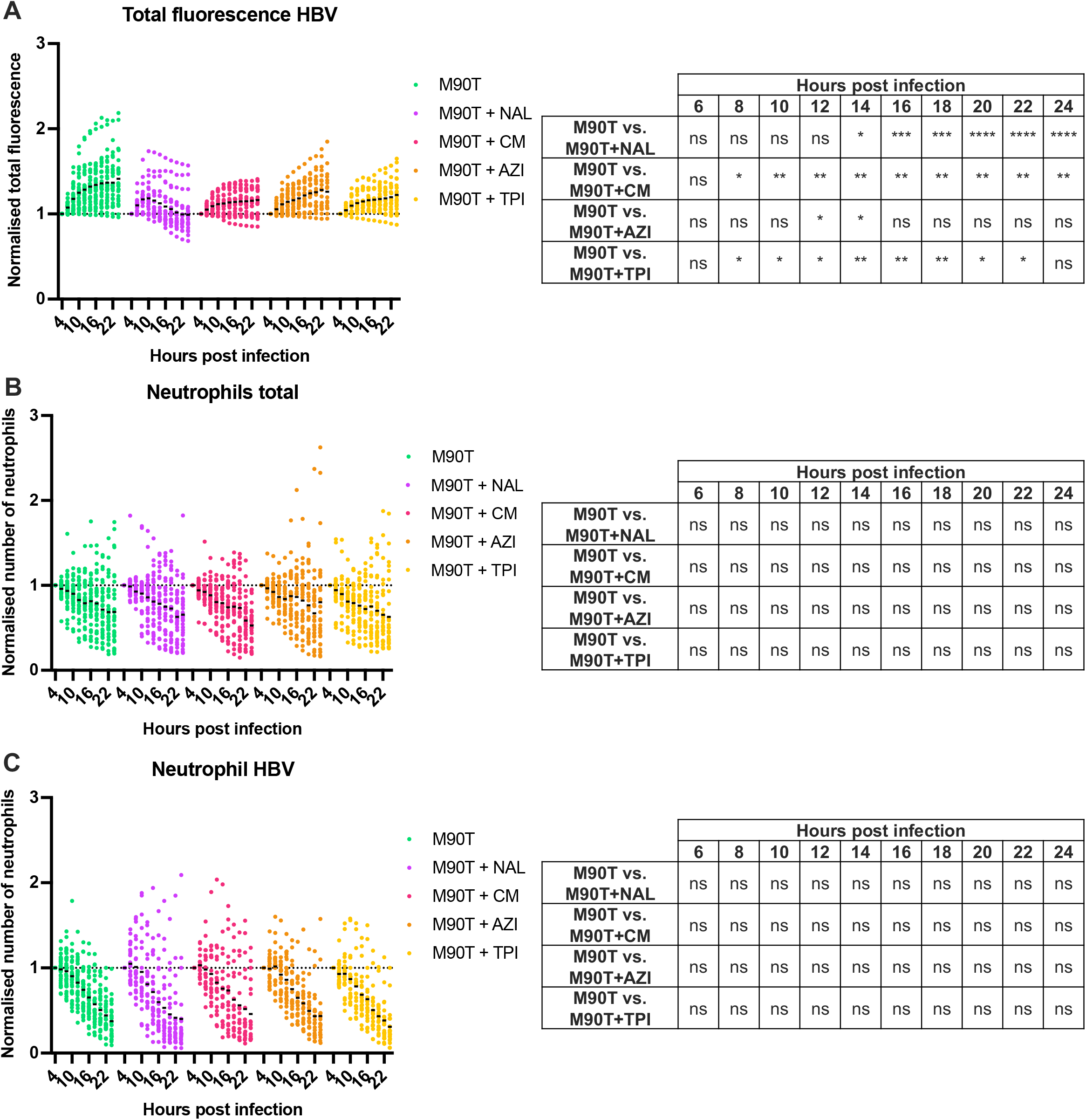
The impact of antibiotics on bacteria and neutrophils during infection. All data presented here is collected from *S. flexneri* M90T-infected 2 dpf zebrafish larvae, non-treated (green) and treated with Nalidixic Acid (NAL, purple), Chloramphenicol (CM, pink), Azithromycin (AZI, orange), or Trimethoprim (TPI, yellow). (A) Normalised total bacterial fluorescence in the HBV. Data normalized to the first timepoint (4 hpi). Data pooled from 3 independent experiments using n > 10 larvae per condition per experiment (ns: non-significant, *p < 0.05, **p < 0.01, ***p < 0.001, ****p < 0.0001. Two-way ANOVA with Dunnett’s multiple comparisons test). (B) Normalised neutrophil quantification at the whole larvae level. Data normalized to the first timepoint (4 hpi). Data pooled from 3 independent experiments using n > 10 larvae per condition per experiment (ns: non-significant. Two-way ANOVA with Dunnett’s multiple comparisons test). (C) Normalised neutrophil quantification in the HBV. Data normalized to the first timepoint (4 hpi). Data pooled from 3 independent experiments using n > 10 larvae per condition per experiment (ns: non-significant. Two-way ANOVA with Dunnett’s multiple comparisons test).

*S. flexneri* infection has been shown to reduce the neutrophil population at the whole zebrafish level (Willis et al., 2018). Treatment with antibiotics did not significantly impact the neutrophil population during infection with either *S. flexneri* or *S. sonnei* (*i*.*e*. neutrophil population was reduced in all conditions, **Fig. 3B, Fig. S3B**). Similarly, treatment with antibiotics did not significantly impact neutrophil recruitment to the HBV (**Fig. 3C, Fig. S3C**).

### Nalidixic acid inhibits bacterial dissemination from the HBV

To further test the impact of antibiotics on infection control *in vivo*, we investigated the dissemination of bacteria from the HBV. In the case of zebrafish HBV infection, we define events of dissemination when *Shigella* is able to cross the blood-brain barrier and spread in the spinal cord. The mechanisms underlying *Shigella* dissemination in zebrafish are mostly unknown, and studying the impact of antibiotics on this process using zebrafish may provide important clues about the control of dissemination in humans.

While dissemination from the HBV in zebrafish is frequent in *S. sonnei* infection (∼ 75%, **Fig. 4A, 4B**), in *S. flexneri* infection it is significantly less frequent (∼ 15%, **Fig. 4C, 4D**). Strikingly, NAL treatment significantly reduced the occurrence of dissemination events to 0% in *S. sonnei* infected larvae and to ∼2% in *S. flexneri* infected larvae (**Fig. 4B, 4D**). These results suggest that NAL works synergistically with the immune system to contain infection in the HBV.

**Figure 4:**
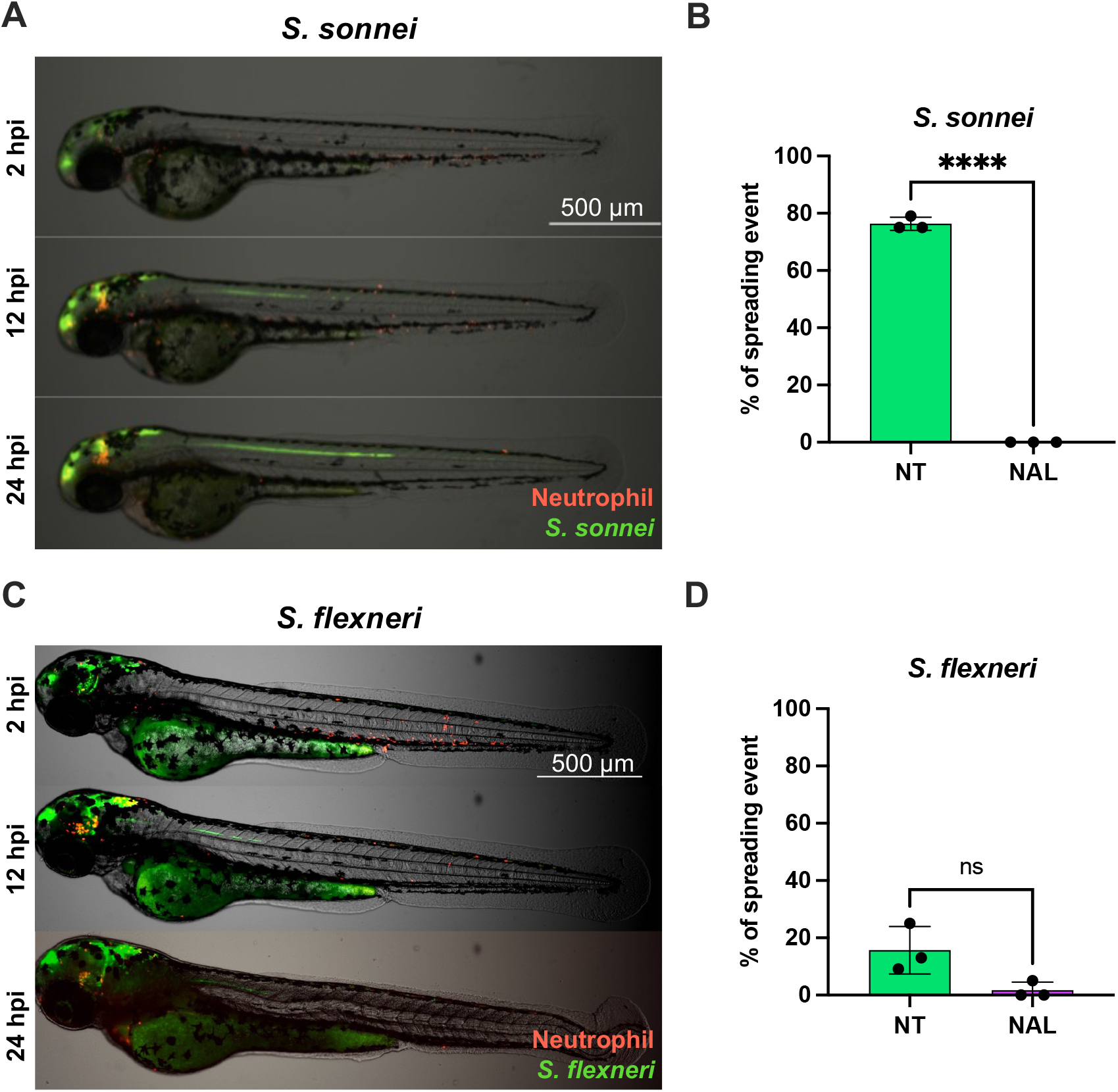
Nalidixic acid prevents *Shigella* dissemination in zebrafish larvae. All data presented here is collected from *S. sonnei* 53G- or *S. flexneri* M90T-infected 2 dpf zebrafish larvae not-treated (green) or treated with Nalidixic Acid (NAL, purple). (A) Representative spreading event in larva infected with *S. sonnei* 53G (green), with labelled neutrophils (red), imaged at 2, 12 and 24 hpi. The spreading event is seen as the green line elongating in the larvae spine. Scale bar: 500 μm. (B) Percentage of spreading events in larvae infected with *S. sonnei* 53G over the course of 24 hpi. Data pooled from 3 independent experiments using n > 10 larvae per condition per experiment (****p < 0.0001. Unpaired Student’s t-test). (C) Representative spreading event in larva infected with *S. flexneri* M90T (green), with labelled neutrophils (red), imaged at 2, 12 and 24 hpi. The spreading event is seen as the green line elongating in the larvae spine. Scale bar: 500 μm. (D) Percentage of spreading events in larvae infected with *S. flexneri* M90T over the course of 24 hpi. Data pooled from 3 independent experiments using n > 10 larvae per condition per experiment (ns: non-significant. Unpaired Student’s t-test).

## DISCUSSION

The emergence of antibiotic resistance in *Shigella* is a global concern, and solutions will require development of innovative antimicrobial strategies. The use of zebrafish has previously helped to illuminate the crucial role of neutrophils in *Shigella* infection control (Duggan and Mostowy, 2018; Gomes and Mostowy, 2020; Mostowy et al., 2013; Torraca and Mostowy, 2018). In this report, we use high content microscopy and develop an automated workflow to non-invasively measure bacterial burden and neutrophils in infected larvae over time. Our workflow significantly accelerates image analysis, and rapidly provides higher throughput of data (e.g. 96 well plates). Since our workflow is flexible, it can be adapted to a large variety of fluorescently labelled bacterial species (as long as the standard curve comparing total fluorescence and CFUs is generated for each species under investigation) or zebrafish immune cells (such as macrophages or hematopoietic stem cells).

Our results using zebrafish show that antibiotic MICs are different *in vitro* and *in vivo*. This may be due to mechanisms of antibiotic storage by specific compartments of the cells (such as lipid droplets), followed by a controlled release of the drugs in the organism (Greenwood et al., 2019). Furthermore, our results indicate that antibiotics are working in synergy with the immune system to control bacterial infection. These results using antibiotics can be compared to results using predatory bacteria (*Bdellovibrio bacteriovorus)* as a treatment against *S. flexneri* infection in zebrafish (Willis et al., 2016). In this case, *Bdellovibrio* injection triggers neutrophil recruitment and promotes direct interaction of *Bdellovibrio* with *S. flexneri*, while antibiotics (diluted into the bath water) require time to diffuse through the larvae to reach the bacteria. It is thus of great interest to precisely compare the mechanism and impact of protection offered by antibiotics versus *Bdellovibrio* using our *Shigella-*zebrafish infection model.

## Concluding remarks

In the future, our automated microscopy workflow can help screen drug libraries, study other clinically relevant *Shigella* strains, and test the role of host genes underlying neutrophil biology. It also has great potential to define the efficacy and MICs of antibiotics *in vivo*. In this way, our workflow may help to define new therapeutic strategies to combat *Shigella* infection in humans. Our workflow can be viewed as an important step towards the development of personalised medicine, and may one day be useful to generate a database of antibiotics and their efficacies against different bacterial pathogens *in vivo*.

## MATERIALS AND METHODS

### Ethics statement

All animal experiments were conducted in concordance with the Home Office and performed on non-protected larvae no older than 4 days post-fertilization (dpf), in accordance with the Animals (Scientific Procedures) Act 1986.

### Bacterial strains and zebrafish lineages

*Shigella flexneri* M90T and *Shigella sonnei* 53G strains both carrying a GFP expressing plasmid (containing a carbenicillin resistance cassette) were used (Mostowy et al., 2010; Valdivia et al., 2006). Bacteria were systematically grown at 37°C in Tryptic Soy Broth (TSB) or on Tryptic Soy Agar (TSA) plates including 0.01% Congo Red to select for clones with a functional T3SS system. The growth medium was supplemented with 100 μg/ml carbenicillin.

Outbred WT zebrafish larvae and the neutrophil transgenic line Tg(*lyz*::DsRed)^*nz50*^ (Hall et al., 2007) were used. Eggs were obtained from naturally spawning zebrafish. They were kept at 28.5°C in 0.5X E2 medium and exposed to 14h of daylight. For experiments involving Phenylthiourea (PTU) treated larvae, embryos were incubated in E2 media + 0.2 mM PTU for 24 h prior to injections (Karlsson et al., 2001).

### Infection assays

Infection assays were performed as previously described (Willis et al., 2018). For infection experiments, individual colonies were grown overnight in 5 ml TSB + 100 μg/ml carbenicillin at 37°C, 200 rotations per minute (rpm). To obtain exponentially grown bacteria (optical density at 600nm (OD600) ∼ 0.6), 400 μl of overnight culture were diluted in 20 ml of TSB + 100 μg/ml carbenicillin media and grown for 2h for a M90T culture, or 1h30 for a 53G culture. The bacterial subculture was then centrifuged for 4 min at 4000g and room temperature (RT). The obtained bacterial pellet was resuspended in 1ml 1X PBS and centrifuged 1 min at 6000g. The pellet was resuspended in 400 μl PBS, and the OD600 of a 50X diluted (in PBS) sample of the bacterial suspension was measured.

The OD600 of the suspension was then adjusted in 50/50 volumes of 0.5% phenol red and 4% Polyvinylpyrrolidone (PVP) in order to reach OD 5 (*i*.*e*. 2 500 Colony Forming Units (CFUs)) or OD 40 (*i*.*e. >* 10 000 CFUs).

Before injection, 2 dpf larvae were prepared and dechorionated if necessary. They were anesthetised in 0.4 mg/ml (2X) tricaine. 5 μl of the suspension to inject (PBS or bacterial suspension) was loaded into a glass capillary needle, which was manually opened to inject 1 nl or 2 nl in the larvae hindbrain ventricle (HBV). The injection was performed with a pressure of nitrogen (N2) between 35 and 40 psi, for 200 milliseconds.

### Quantification of bacterial burden

To quantify and characterize the bacterial inoculate, 3 injected larvae per condition were transferred into microcentrifuge tubes individually containing 200 μl of 0.1% Triton X-100 diluted in PBS. The larvae were then mechanically disrupted and dissolved in the solution using sterile pestles. The suspensions were serially diluted and 20 μl were plated on TSA plates supplemented with 100 μg/ml carbenicillin and 0.01% Congo Red, and then incubated for 18h at 37°C. CFUs were hand counted the next day to assess the bacterial burden in infected larvae.

### MICs determination and Antibiotic treatment assays

To assess the minimum inhibitory concentrations (MICs) for every antibiotic tested on the 2 strains (*S. flexneri* M90T and *S. sonnei* 53G), antibiotic containing stock solutions were prepared at 8 μg/ml for Nalidixic acid (NAL), 160 μg/ml for Chloramphenicol (CM), 16 μg/ml for Azithromycin (AZI) and 2 μg/ml for Trimethoprim (TPI). These solutions were then serially diluted 5 times in TSB, to generate 5 daughter solutions for each antibiotic. In detail, NAL solutions were at 4, 2, 1, 0.5, 0.25 and 0.125 μg/ml, CM at 80, 40, 20, 10, 5 and 2.5 μg/ml, TPI at 1, 0.5, 0.25, 0.125, 0.063 and 0.032 μg.ml and AZI at 8, 4, 2, 1, 0.5 and 0.25 μg/ml. 200 μl of every solution was transferred in the wells of several 96 well plates. 1 μl of overnight bacterial culture was used to inoculate the wells for every condition, and standard wells containing 200 μl of TSB only. Every condition was tested in triplicate. The plates were sealed and incubated at 37°C, 200 rpm for 7 hours. The growth was then assessed visually for every condition. MICs were considered as the concentrations in the first well without noticeable growth. Importantly, only the MICs for 53G (the highest concentrations) were considered for antibiotic treatment assays.

To test the effect of drugs at these concentrations, 10 larvae were grown for 24 hours in E2 with the corresponding antibiotic at the *S. sonnei* MIC, in a 6 well plate. Developmental defects were evaluated by eye under a binocular microscope.

For *in vivo* antibiotic treatment assays, larvae injected with 10 000 CFUs of *Shigella* were kept for 30 min-1 hour in antibiotics diluted in E2 water and 0.2 mg/mL (1X) tricaine to reach the MIC of the considered drug. This first antibiotic treatment was performed while the rest of the experimental setup was prepared. When ready, the larvae were placed in a 96 well plate (CellCarrier-96 Ultra - PerkinElmer) and embedded in 1% low melting-point agarose (diluted in 0.5X E2 media) to be immobilized. The larvae were meticulously disposed in the wells to increase imaging efficiency: for whole fish imaging they were set horizontally and on the lateral side, and for HBV imaging they were set head down against the glass bottom of the 96 well plate. Once the agarose was set, wells were topped up by antibiotic solutions diluted in E2 + 0.2 mg/mL (1X) tricaine to reach 2X times the MIC of the appropriate antibiotic. This concentration is necessary to observe the same effect of the antibiotic as in free swimming larvae, since agarose interferes with diffusion of the drug (data not shown).

### Microscopy acquisition

Larvae were imaged *in vivo*, while embedded and anesthetised, in a 96-well plate (CellCarrier-96 Ultra - PerkinElmer), using Cell Discoverer 7 (CD7) microscope from Zeiss. For time lapse experiments, images were taken every 2h until 24 hours post-infection (hpi) and the CD7 was heated to 31°C. Whole zebrafish imaging was performed with a 5X/0.35 plan-apochromat objective, and a 0.5X tubelens, and HBV imaging was performed with the same objective but with a 2X tubelens to capture 51 slice Z-stacks over a range of 250 μm. To compare the performance of the ‘Ellett and Lieschke’ method in images acquired with the CD7 microscope and a Leica M205FA stereomicroscope, larvae were imaged using a Leica M205FA stereomicroscope at a magnification of 0.3X.

### Image analysis

Custom macros were designed in Fiji (https://fiji.sc) to automate image analysis. All macros are dependent on information (including folder containing the files to analyse, channel to analyse, parameters to consider, type of analysis desired) provided by the user on a user-friendly interface.

#### Bacterial burden quantification

The bacterial burden quantification macro was designed to analyse Z-stacks of larvae HBV, although it can be adapted to many other image types, such as whole zebrafish images. The macro automatically performs a sum Z-projection on the images, measures the mean fluorescence as well as the image area, and multiplies the two values to obtain the total fluorescence in the Z-stack. The macro automatically runs through files in batch, and returns a text document containing the titles of the images analysed and the total fluorescence associated to each one of them. This total fluorescence value is considered as a proxy of the bacterial burden.

#### Neutrophil detection and characterisation (LenCell)

Neutrophil detection and characterisation can be performed by two different macros depending on the image type to analyse (single plane, or Z stack). In brief, the macro for single plane images automatically performs a bandpass filter with the « subtract background » feature of Fiji. This feature uses the « rolling ball method » (https://imagej.net/plugins/rolling-ball-background-subtraction), which iteratively determines local background values for every pixel in the image by averaging the values over a large ball around the pixel. This local background value is then subtracted from the image. The radius of the rolling ball (RBA) can be changed directly in Fiji and in the macro. To perform the bandpass filter, the macro duplicates two times the original image and subtracts the backgrounds of each of these duplicates using different RBA values. Typically, a large RBA value is chosen (here 11) for one duplicate, to remove large background spots (e.g. yolk sack, skin, contaminations in the agarose), and a small one (here 3) is chosen for the other duplicate, to remove the « small » background, especially at neutrophil clusters. Finally, to retrieve a cleaned result image close to the original one, the pixel values of both duplicates are multiplied. The image obtained after this operation is automatically converted to a binary mask, using a default threshold. This threshold can be modified for other applications. The binary mask is extensively cleaned with a step of opening (erosion + dilation of the black spots), which removes small noise. The resulting image is then processed through a step of watershed (https://imagej.net/plugins/classic-watershed), which automatically detects clusters of black spots and divides them.

The cleaned binary image is finally processed by the « analyse particle » tool of Fiji, which automatically detects objects in the binary mask above the threshold, or regions of interest (ROIs), in the image, and stores information about their coordinates, shape and size. A size filter set is applied (here to 0-infinity), and the number of ROIs is automatically counted, while their boundaries are overlaid on the original image. This provides a visual output to check for any artifact. Additionally, a compressed file containing the positional information of the ROIs detected is automatically saved on the computer.

These steps are entirely automated, although the user must provide the value of the wanted parameters (the 2 RBAs to use, and the minimum size of the size filter) as well as the path to the folder containing the files to analyse. After processing, a table containing the titles of the analysed images and the number of ROIs associated is returned, as well as a series of visual outputs.

This macro can additionally be automatically optimised. To do so, the user hand counts a series of images and provides data to the macro. After asking the range of parameters (RBA values and size filter) that the user wants to train the macro on, it will automatically and iteratively use every combination of parameters to count neutrophils in the provided images. The parameters which provide results closest to the hand count are then considered as ‘optimized parameters’.

The macro to count and detect neutrophils in an HBV Z-stack image is very similar, however it includes several pre-processing steps. First the image is cleaned by using a very large band pass filter (first RBA of 100, second RBA of 200). The two resulting images are added to each other, and the resulting image is processed through a Z projection by standard deviation, which makes the ROIs stand out. The Z projection is finally processed through the same pipeline as described before for the whole zebrafish images, but with slightly differing parameters (first RBA: 9, second RBA: 10, minimum size 0).

### Statistical analysis and data processing

Statistical significance was determined using GraphPad Prism v9 (https://www.graphpad.com/scientific-software/prism/). For comparisons of the different analysis methods, a two-way ANOVA with *post-hoc* Sidak’s multiple comparisons test was performed. For experiments occurring over time, ie. bacterial burden and neutrophil counts over time, are presented as normalised to the first time point for every larvae (4 hpi). A two-way ANOVA with *post-hoc* Dunnett’s multiple comparisons test was performed for these experiments. For quantification of spreading events, an unpaired Student’s t-test was used. Outlier larvae (dead, uninfected) were removed from the dataset. The last timepoint thus typically contains less replicates, as some larvae died overtime. In the figures, p-values are described as: p-value > 0.05: non-significant (ns); p-value < 0.05: *; p-value < 0.01: **; p-value < 0.001: ***; p-value < 0.0001: ****.

## ACKNOWLEDGEMENTS

We thank all members of the Mostowy lab and Zeiss engineers for helpful discussions.

## DATA AVAILABILITY

Datasets will be provided by the corresponding authors upon request. LenCell source code is available under https://github.com/arthurlensen/LenCell.git.

## AUTHORS CONTRIBUTION STATEMENT

Conceptualization: A.L., M.C.G, A.T.L.J., S.M.; Methodology: A.L., M.C.G, A.T.L.J.; Software: A.L., M.C.G, A.T.L.J.; Formal analysis: A.L., M.C.G, A.T.L.J.; Resources: A.L., M.C.G, A.T.L.J., S.M.; Data curation: A.L., M.C.G, A.T.L.J., S.M.; Writing: A.L., M.C.G, A.T.L.J., S.M.; Supervision: M.C.G, A.T.L.J., S.M.; Funding acquisition: S.M.

